# Cerebral activity in monkeys *Papio anubis* during the perception of conspecific and heterospecific agonistic vocalizations: A functional Near Infrared Spectroscopy study

**DOI:** 10.1101/2022.04.21.489037

**Authors:** Coralie Debracque, Thibaud Gruber, Romain Lacoste, Adrien Meguerditchian, Didier Grandjean

## Abstract

Are conspecific emotional vocalizations special? Although often investigated in non-human primates using functional magnetic resonance imaging or positron emission tomography, it remains unclear whether the listening of conspecific vocal emotions leads to similar or different cerebral activations when compared to heterospecific calls (i.e. expressed by another primate species). Using a neuroimaging technique rarely employed in monkeys so far, functional Near Infrared Spectroscopy (fNIRS), the present study investigated cortical temporal activities during exposure to both conspecific and heterospecific calls in three female adult baboons (*Papio anubis*). The three subjects were lightly anesthetized and passively exposed to agonistic baboon and chimpanzee (*Pan troglodytes*) vocalizations, as well as energy matched white noises in order to control for this low-level acoustic feature. Despite inter-individual variabilities, permutation test analyses on the extracted OxyHemoglobin signal revealed for two subjects out of three significant differences between the passive listening of baboon *versus* chimpanzee stimuli. Additionally, in one subject, a modulation of the left temporal cortex activity was found for the perception of baboon calls contrasted to chimpanzee vocalizations as well as for the passive listening of baboon white noises compared to chimpanzee ones. Although the lack of generalization of those findings in all three subjects prevents us to drawn any conclusion and that more subjects would be needed, the hypothesis that baboons’ cortical temporal regions may be more sensitive to the processing of conspecific sounds compared to heterospecific stimuli is not excluded. Our study highlights that fNIRS may be a promising alternative to further investigate the auditory mechanisms at play in the right and left baboons’ temporal cortices for the processing of emotional vocalizations.

## 1. Introduction

Since the 1990s (George et al. 1995; Pihan, Altenmüller, and Ackermann 1997), many neuroimaging studies have investigated the human temporal cortex activity in emotional voice processing. Hence, functional Magnetic Resonance Imaging (fMRI) and more recently functional Near Infrared Spectroscopy (fNIRS) data pointed out the role of the bilateral superior temporal gyrus (STG), superior temporal sulcus (STS) and middle temporal gyrus (MTG) in the processing of emotional prosody (Grandjean 2020; Grandjean et al. 2005, 2005; Kotz et al. 2013; Plichta et al. 2011; Wildgruber et al. 2009; Zatorre and Belin 2001) and more specifically in the recognition of positive and negative emotions (Bach et al. 2008; Frühholz and Grandjean 2013; Johnstone et al. 2006; Zhang, Zhou, and Yuan 2018).

Yet, despite calls for an evolutionary-based approach to emotions that consider the adaptive functions and the phylogenetic continuity of emotional expression and identification (Bryant 2021; Greenberg 2002), few comparative research investigated the human temporal cortex activity during the recognition of emotional cues in human voices (conspecific) and other species vocalizations (heterospecific), expressed especially by non-human primates (NHP), our closest relative (Perelman et al. 2011). In the few studies published so far, fMRI analysis has revealed more activations in human auditory cortex for the recognition of agonistic vocalizations expressed by humans as well as macaques (*Macaca mulatta*) and domestic cats (*Felis catus*) comparing to affiliative ones. This result was largely driven by a decrease of activity in auditory cortex during the listening of macaque and cat vocalizations compared to human vocalizations (Belin et al. 2008). Additionally, Fritz and colleagues demonstrated a greater involvement of the human STS and the right planum temporale (PT) for the identification of human emotional voices contrasted to chimpanzee (*Pan troglodytes*) and then macaques calls (Fritz et al. 2018). Interestingly, similar results were found in the STS and STG when human emotional voices were compared to various animal sounds, non-vocal stimuli or non-biological noises (Bodin et al. 2021; Pernet et al. 2015) suggesting a specific sensitivity of the superior regions of the human temporal cortex for conspecific voices.

Is this sensitivity of the temporal cortex to emotional cues expressed by conspecific found in NHP? In other words, are the cerebral mechanisms of vocal emotion perception shared across primate species, or has the auditory cortex of *Homo sapiens* evolved differently?

The previous literature on primates emphasizes the brain continuity between humans and NHP for the auditory processing of conspecific emotions. For instance, fMRI studies in macaques have revealed a greater involvement of the STG for the perception of conspecific emotional calls compared to heterospecific ones including calls from other primate and animal species, environmental sounds and scrambled vocalizations (Joly et al. 2012; Ortiz-Rios et al. 2015; Petkov et al. 2008). Following this, positron emission tomography (PET scan) studies have shown the predominant role of the right PT in chimpanzees (Taglialatela et al. 2009) and of the STS in macaques (Gil-da-Costa et al. 2004) for the processing of conspecific emotional calls. Additionally, neurobiological findings in macaques and marmosets (*Callitrix jacchus*) have suggested a greater involvement of the STG and of the primary auditory cortex in the passive listening of emotional conspecific calls comparing to environmental sounds, scrambled or time-reversed vocalizations (Belin 2006; Ghazanfar and Hauser 2001; Poremba et al. 2004). Despite these results, the question on the specific status of conspecific emotions in NHP remains poorly explored with respect to heterospecific vocalizations. In particular, because of the species-dependent results in humans highlighted above and the phylogenetic proximity across primate species, it seems necessary to include heterospecific stimuli from other NHP to reconstruct the phylogenetic evolution of primate vocal emotion processing (Bryant 2021).

The present study investigated temporal cortex involvement in three female baboons: Talma, Rubis and Chet, during exposure to conspecific vs. heterospecific affective vocalizations, using fNIRS. Building on a growing interest over the past decade (Boas et al. 2014; Pan, Borragán, and Peigneux 2019), we used fNIRS because of its non-invasiveness, its poor sensitivity to motion artefacts (Balardin et al. 2017) and its suitability for comparative research (Debracque et al. 2021; Fuster et al. 2005; Kim et al. 2017; Lee et al. 2017; Wakita et al. 2010). According to the existing literature on NHP and humans suggesting a sensitivity of the primates’ temporal cortex for conspecific calls, we expected *i)* more activation in the temporal cortex for the passive listening of baboon sounds compared to chimpanzee stimuli; and *ii)* a greater involvement of the temporal cortex for the perception of agonistic conspecific vocalizations in comparison to the other sounds.

## 2. Material & Methods

### 2.1. Subjects

Three healthy female baboons (Talma, Rubis and Chet, mean age = 14.6 years, SD ± 3.5 years) were included in the present study. Based on the annual health assessment and the daily behavioural surveys made by the veterinary and animal welfare staff, the subjects had normal hearing abilities and did not present a structural neurological impairment (confirmed with respective T1w anatomical brain images – 0.7 × 0.7 × 0.7 resolution – collected in vivo under anesthesia in the 3Tesla MRI Brunker machine). All procedures were approved by the “C2EA -71 Ethical Committee of neurosciences” (INT Marseille) and performed in accordance with the relevant French law, CNRS guidelines and the European Union regulations. The subjects were born in captivity and housed in social groups at the Station de Primatologie in which they have free access to both outdoor and indoor areas. All enclosures are enriched by wooden and metallic climbing structures as well as substrate on the group to favour foraging behaviours. Water is available *ad libitum* and monkey pellets, seeds, fresh fruits and vegetables were given every day. The three subjects were lightly anesthetized with propofol and passively exposed to auditory stimuli as described below (see also Debracque et al. 2021 for the complete protocol).

### 2.2. Stimuli

Auditory stimulations consisted of agonistic vocalizations expressed by baboon (conspecific – see Figure 1a) and chimpanzee (heterospecific – see Figure 1b) species as well as energy matched white noises to control for this low-level acoustic feature and for its unfolding (i.e. the temporal structure of energy of the vocalizations). Each stimulus had a duration of 20 seconds, and was repeated six times (see Debracque et al. 2021 for more details). The auditory stimuli were pseudo-randomized, alternating vocalizations and white noises; and were separated by 15 seconds of silence. Additionally, auditory stimulations were broadcasted either binaurally or monaurally in the right or left ear.

**Figure 1:**
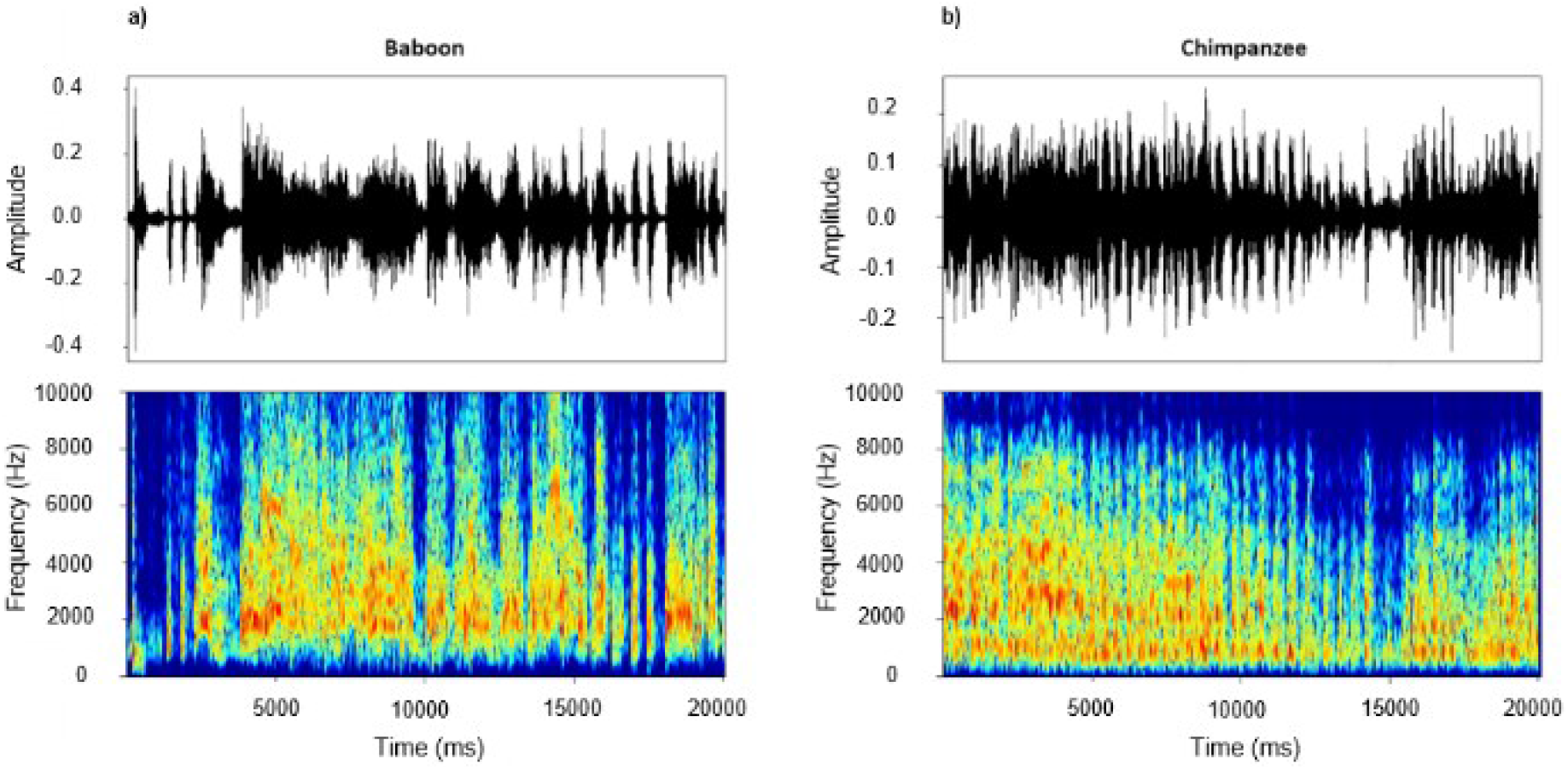
Representative waveforms and spectrograms of 20s-long agonistic **a)** baboon and **b)** chimpanzee vocalizations used as stimuli in the present study. These graphical representations were extracted using the PhonTools package (Barreda 2015) on R. studio (Team 2020).

### 2.3. fNIRS

#### 2.3.1. Recordings

Brain activations were measured using two light and wireless fNIRS devices (Portalite, Artinis Medical Systems B.V., Elst, The Netherlands) enabling the measurement of Oxyhemoglobin (O_2_Hb). Optical probes were placed on the right and left temporal cortices of the subjects using T1 MRI scanner images previously acquired at the Station de Primatologie on baboons (see Figure 2). Data were obtained at 50 Hz with two wavelengths (760 and 850 nm) using three channels per hemisphere (ch1, ch2, ch3) with three inter-distance probes (3 – 3.5 – 4 cm) investigating three different cortical depths (1.5 – 1.7 – 2 cm respectively).

**Figure 2:**
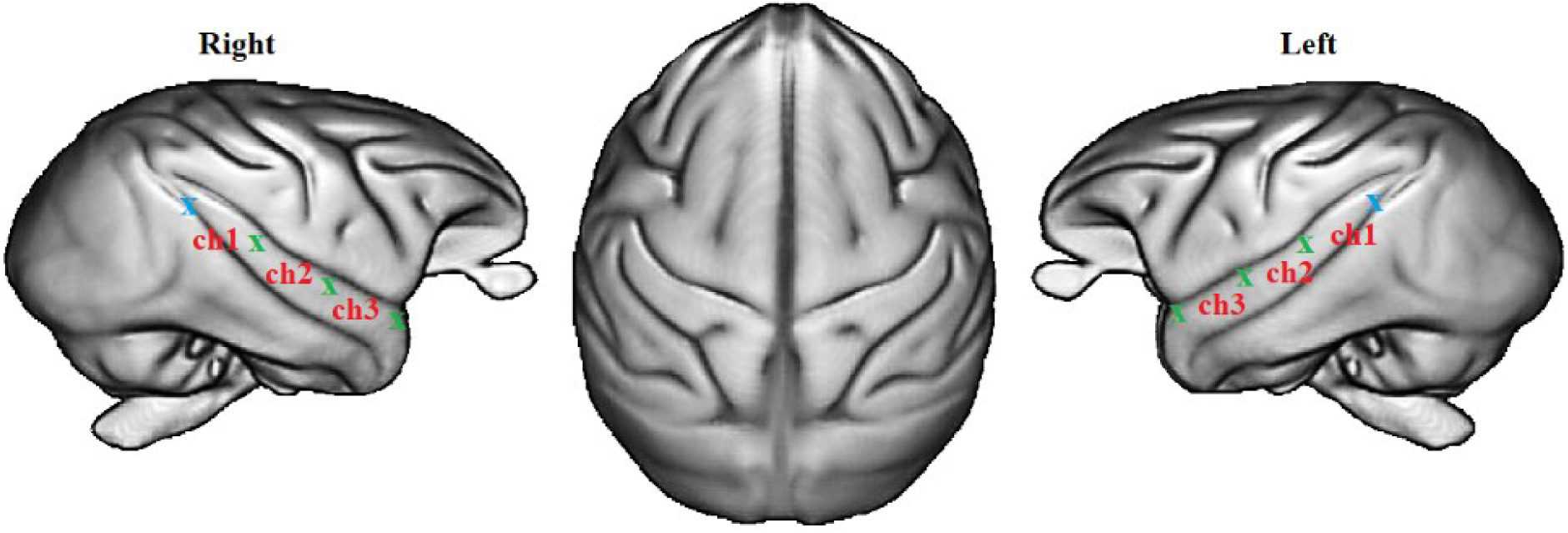
fNIRS optode and channel locations according to 89 baboons T1 MRI template (Love et al. 2016). Blue and green crosses represent optical receivers and transmitters respectively. Ch1, Ch2 and Ch3 indicate the three channels on the right and left temporal cortex.

Reducing the potential disturbing impact of the fNIRS protocol on the subjects, each experimental session was planned during the baboons’ routine health inspection at the Station de Primatologie. As part of the health check, subjects were isolated from their social group and anesthetized with an intramuscular injection of ketamine (5 mg/kg - Ketamine 1000®) and medetomidine (50μg/kg - Domitor®). Then Sevoflurane (Sevotek®) at 3 to 5% and atipamezole (250 μg/kg - Antisedan®) were administered before recordings. Each baboon was placed in ventral decubitus position on the table and the head of the individual was maintained using foam positioners, cushions and Velcro strips to remain straight and to reduce potential motion occurrences. Vital functions were monitored (SpO2, Respiratory rate, ECG, EtCO2, T°) and a drip of NaCl was put in place during the entire anaesthesia. Before fNIRS recordings, temporal areas on the baboons’ scalp were shaved and sevoflurane inhalation was stopped. Subjects were further sedated with a minimal amount of intravenous injection of Propofol (Propovet®) with a bolus of around 2mg/kg every 10 to 15 minutes or by infusion rate of 0.1 – 0.4 mg/kg/min. After the recovery period, subjects were put back in their social group and monitored by the veterinary staff.

#### 2.3.2. Analysis

SPM_fNIRS toolbox (Tak et al. 2016) and custom made codes on Matlab 7.4 R2009b (The MathWorks Inc. 2009) were used to perform first level analysis on raw fNIRS data following this procedure: *i)* O_2_Hb concentration changes were calculated using the Beer-Lambert law (Delpy et al. 1988); *ii)* motion artifacts were removed manually in each individual and each channel. In total, 10 seconds (1.3%) were removed from the O_2_Hb signal of Rubis and 35 seconds (4.8%) for Talma and Chet; *iii)* a low-pass filter based on the HRF (Friston et al. 2000) was applied to reduce physiological confounds; *iv)* a baseline correction was applied in subtracting the pre-stimulus baseline from the post-stimulus O_2_Hb concentration changes of each trial and *v)* O_2_Hb signal was averaged for Talma in a window of 4 to 12 s post stimulus onset for each trial; and for Rubis and Chet in a window of 2 to 8 s post stimulus onset in order to select the range of maximum O_2_Hb concentration changes following Debracque et al 2021. Shortly, the differences of concentration range are explained by the presence of tachycardia episodes for both Rubis and Chet during the experiment, involving an HRF almost twice as fast as the one found for Talma.

The second level analysis was made on R. studio (Team 2020) using the permuco package (Frossard and Renaud 2019). In each Hemisphere (right, left), we used non-parametric permutation tests with 5000 iterations to assess O_2_Hb concentration changes for each subject (Talma, Rubis, Chet) as they enable repeated measures ANOVA in small sample sizes by multiplying the design and response variables (Kherad-Pajouh and Renaud 2015). *Stimuli* (call, white noise); *Species* (baboon, chimpanzee); *Channels* (ch1, ch2, ch3) and *Sides* (right, left, both ears) were selected as fixed factors. As recommended, contrast effects of *Species* and *Stimuli* within channels were assessed with 2000 permutations (Kherad-Pajouh and Renaud 2015). Both p. values for permutation *F* (p_perm_) and parametric *F* are reported.

## 3. Results

The main effect *Channels* was significant in nearly all subjects and hemispheres: Talma (right: *F*(2,3) = 161, 5, p | p_perm_ <.001; left: *F*(2,3) = 33.91, p | p_perm_ <.001); Rubis (right: *F*(2,3) = 8, 99, p | p_perm_ <.001); and Chet (right: *F*(2,3) = 3,99, p | p_perm_ <.05; left: *F*(2,3) = 25.68, p | p_perm_ <.001). It was not significant for Rubis’ left hemisphere (*F*(2,3) = 2.15, p | p_perm_ =.12). Hence, we reported the effects of *Species* and *Stimuli* according to channels in Figure 3. Note that for all subjects, the factor *Sides* did not reach significance and thus, do not statistically explain differences for *Stimuli* and *Species* (see Debracque et al. 2021 for more details related to auditory asymmetries). All results are reported in Supplementary Materials.

**Figure 3:**
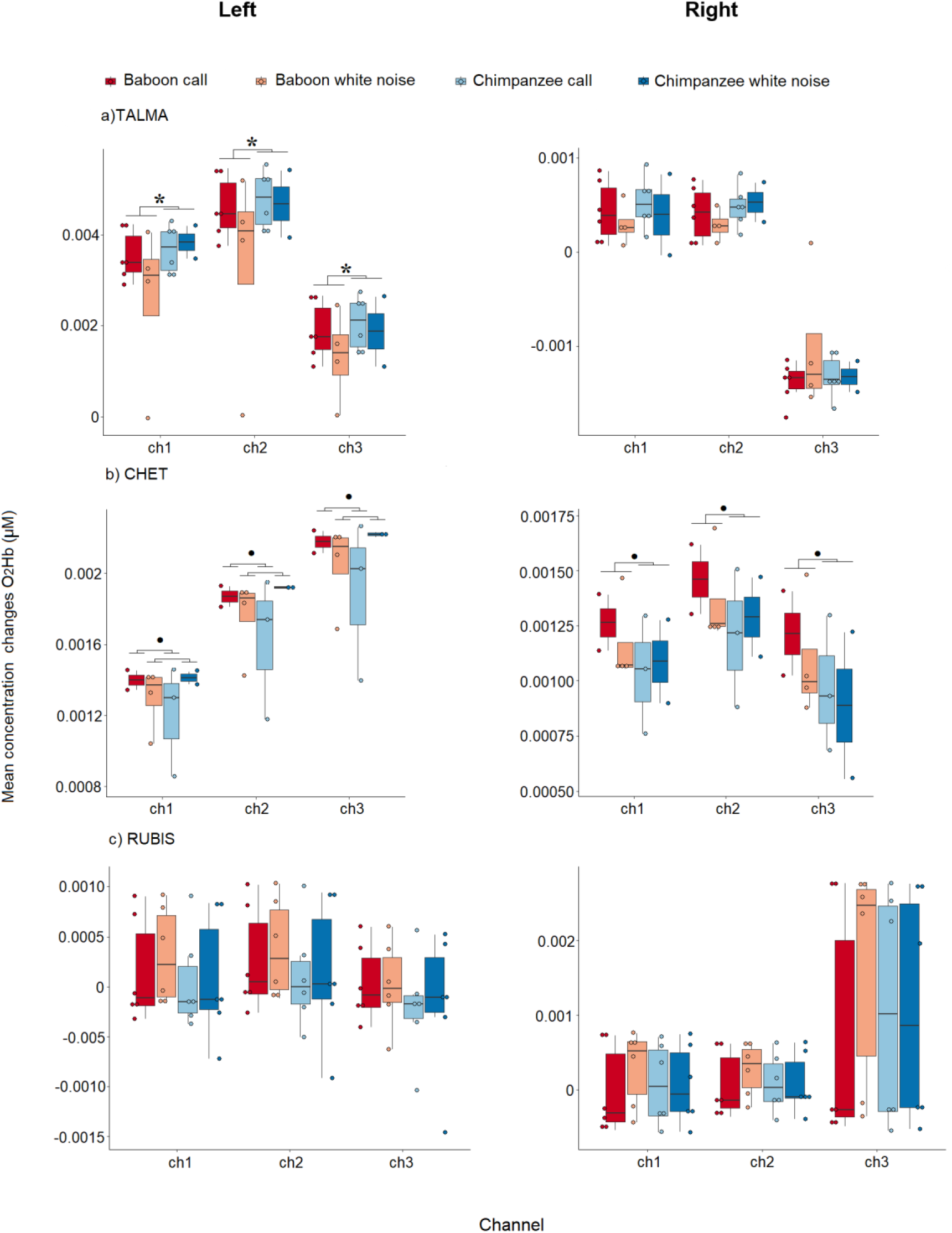
Right and left temporal cortex activations for the baboons **a)** Talma, **b)** Chet and **c)** Rubis during the perception of agonistic baboon (conspecific) and chimpanzee (heterospecific) vocalizations as well as their energy matched white noises. The mean concentration changes of O_2_Hb (*y* axis) is represented in micro molar (μM) for each fNIRS channel (*x* axis). Colourful dots and dark lines represent stimuli and confidence intervals respectively. Results of the permutation tests within channels are shown with **\*** p<.05; p=.07. The ggplot2 package (Wickham et al. 2021) on R.studio (Team 2020) was used for visualizing the data.

Regarding the right hemisphere, permutation tests revealed for Chet a significant main effect of the factor *Species* (*F*(1,2) = 5.03, p | p_perm_ <.05).

Regarding the left hemisphere, the main factors *Species* was found significant for the baboon Talma (*F*(1,2) = 4.24, p| p_perm_ <.05). In addition, a significant two-way interaction was found significant for the subject Chet between *Stimuli* and *Species* (*F*(1,4) = 4,13, p | p_perm_ =.05).

In sum, across the three channels, more O_2_Hb concentration changes were found in Talma’s left hemisphere for the perception of chimpanzee sounds compared to baboon stimuli. Conversely, for Chet, permutation analyses revealed more O_2_Hb concentration changes in the right hemisphere for the passive listening of baboon sounds comparing to chimpanzee ones. Additionally, her left hemisphere was more activated by baboon calls vs. chimpanzee calls and by chimpanzee white noises vs. baboon white noises.

## 4. Discussion

The present fNIRS study suggests, in two out of our three tested baboons, a sensitivity of the baboons’ temporal cortex for conspecific sounds with documented differential activations in contrast to heterospecific sounds.

This finding is quite consistent with the few existing fMRI data in macaques showing the distinctive role of STG in the perception of emotions in conspecific and heterospecific calls (Joly et al. 2012; Petkov et al. 2008). Similar to humans (Belin et al. 2008; Fritz et al. 2018), this first result suggests a different involvement of temporal regions in NHP for the processing of emotions in conspecifics and heterospecific primate vocalizations. From an evolutionary perspective, this would be a mechanism inherited from the common ancestor of *Homo sapiens* and other *Catharrini* around 40 million years ago (Harrison 2013), although it may be earlier if *Platyrrhini* share this feature too.

Despite the significant main effect *Species* for two subjects, fNIRS data revealed also inter-individual differences between Talma and Chet. First, we failed to record significant results for Rubis, although this may have been a consequence of her constant tachycardia during the health check and experiment (see Debracque et al. 2021). Second, the left temporal cortex of Talma was overall more activated for chimpanzee sounds (calls and white noise) than for baboon ones; in contrast, results were reversed in the left temporal cortex of Chet, where statistical analyses highlighted an increase of O_2_Hb concentration changes in the left temporal cortex for the passive listening of agonistic baboon sounds compared to chimpanzee sounds. In addition, in her right temporal cortex, we documented an increase in O_2_Hb concentration led by the perception of agonistic baboon call.

Differences between individuals in temporal cortex sensitivity could explain this absence of congruence in our fNIRS data. Xu and colleagues (2019), have in fact, demonstrated in five anaesthetized and awake macaques great individual variabilities in functional connectivity across cortical regions (Xu et al. 2019). Interestingly, the authors compared macaques’ fMRI data to human ones and concluded to a similar heterogeneity in functional connectivity across primate species. In the same line, Pernet and colleagues also demonstrated, using fMRI, high inter-individuality differences between human STG and STS for the listening of conspecific emotional voices compared to non-vocal sounds (Pernet et al. 2015). Hence, as for highly cited human neuroimaging studies (Szucs and Ioannidis 2020), future non-invasive comparative research should include more subjects to take in consideration this inter-individual variability in brain mechanisms. Yet, the necessity to increase NHP subjects to address limits in statistical power faces ethical aspects related to animal welfare in the case of invasive neuroimaging studies. The development of fNIRS (Debracque et al. 2021) and longitudinal studies in comparative neuroscience (Song et al. 2021) would thus allow answering parts of these challenges.

Often explored using fMRI or Pet scan (e.g. Bodin et al. 2021; Gil-da-Costa et al. 2004; Ortiz-Rios et al. 2015), our fNIRS data remain inconclusive regarding any differences in temporal cortex activity between the passive listening of agonistic baboon vocalizations and white noises. Indeed, comparative research on macaques and marmoset (*Callithrix jacchus*) showed a greater sensitivity of the temporal cortex in its anterior part than in its posterior area for the contrast conspecific calls vs. control sounds (Bodin et al. 2021). Future fNIRS studies should improve the probe location on the scalp of baboons or any other NHP species.

Finally, out of the scope of this paper, permutation tests and descriptive analyses highlighted consistent fNIRS data for the channels 1 and 2 compared to channel 3 on both, right and left hemispheres. This result suggests that, for fNIRS in baboons, the best inter-probe distant to assess cortical activations would be between 3cm and 3.5cm. Interestingly, these distances are commonly used for fNIRS experiments in human adults (Ferrari and Quaresima 2012).

Overall, highlighting the evolutionary continuity between humans and NHP, our results suggest the existence in baboons of differences in temporal cortex activity between the processing of conspecific and heterospecific sounds. Yet, fNIRS data also pointed out high inter-individual variability in these results and remain inconclusive in regards to the contrast conspecific agonistic vocalizations vs. white noises (control sounds), which are often explored meaningfully using fMRI or Pet scan (e.g. Bodin et al. 2021; Gil-da-Costa et al. 2004; Ortiz-Rios et al. 2015). Moreover, the present study has some limitations. Firstly, our experiment focused on baboons and it is unclear whether it will replicate in other monkey species. In fact, Fitch and Braccini (2013) have already suggested differences between monkey in terms of mechanisms for the processing of conspecific and heterospecific vocalizations (Fitch and Braccini 2013). Finally, only agonistic vocalizations were included in the present fNIRS protocol. Yet, similar to humans, NHP might have some distinctive brain mechanisms for negative and positive emotions (e.g. Davidson 1992; Frühholz and Grandjean 2013; Zhang et al. 2018). Consequently, future studies should consider the inclusion of positive emotions, more individuals or the use of fNIRS and longitudinal studies, as well as various monkey species in order to maximize the generalization of their neuroimaging data on auditory processing of emotions in NHP.

## Supporting information

Supp mat

## Acknowledgements

We thank the Swiss National Foundation and the European Research Council for funding this work. We warmly thank the vet Pascaline Boitelle, the animal care staff as well as Jeanne Caron-Guyon, Lola Rivoal, Théophane Piette and Jérémy Roche for assistance during the recordings. CD also thanks Dr. Ben Meuleman for his wise advices on permutation test analyses.

## Declarations

### Availability of data and material

Raw data are freely available to any researcher.

### Code availability

Custom Matlab and R. studio codes are available on request.

### Ethics approval

All animal procedures were approved by the “C2EA -71 Ethical Committee of neurosciences” (INT Marseille) under the application number APAFIS#13553-201802151547729, and were conducted at the Station de Primatologie CNRS (UPS 846, Rousset-Sur-Arc, France) within the agreement number C130877 for conducting experiments on vertebrate animals. All methods were performed in accordance with the relevant French law, CNRS guidelines and the European Union regulations (Directive 2010/63/EU).

### Consent to participate

Not applicable.

### Consent for publication

Not applicable.

