## Supplementary material for "Cerebral activity in monkeys *Papio anubis* during the perception of conspecific and heterospecific agonistic vocalizations: A functional Near Infrared Spectroscopy study": Supp mat

**Table S1:** Results from permutation tests with 5000 permutations for each hemisphere and subject. Abbreviations: (RH) right hemisphere; (LH) left hemisphere. Significant analyses are highlighted in light green.

| | Talma | | Rubis | | Chet | |
|---|---|---|---|---|---|---|
| | **RH** | **LH** | **RH** | **LH** | **RH** | **LH** |
| ***Main effect*** | | | | | | |
| *Species* | $F(1,2)=0.34$ $p\|p_{perm}=.57$ | $F(1,2)=4.24$ $p\|p_{perm}<.05$ | $F(1,2)=0.13$ $p\|p_{perm}=.71$ | $F(1,2)=2.59$ $p\|p_{perm}=.12$ | $F(1,2)=5.03$ $p\|p_{perm}<.05$ | $F(1,2)=0.24$ $p\|p_{perm}=.62$ |
| *Stimuli* | $F(1,2)=0.04$ $p\|p_{perm}=.95$ | $F(1,2)=1.87$ $p\|p_{perm}=.18$ | $F(1,2)=1.21$ $p\|p_{perm}=.27$ | $F(1,2)=0.25$ $p\|p_{perm}=.62$ | $F(1,2)=0.13$ $p\|p_{perm}=.72$ | $F(1,2)=0.79$ $p\|p_{perm}=.38$ |
| *Channels* | $F(2,3)=161$ $p\|p_{perm}<.001$ | $F(2,3)=33.9$ $p\|p_{perm}<.001$ | $F(2,3)=8.99$ $p\|p_{perm}<.001$ | $F(2,3)=2.15$ $p\|p_{perm}=.12$ | $F(2,3)=3.99$ $p\|p_{perm}<.05$ | $F(2,3)=25.7$ $p\|p_{perm}<.001$ |
| *Sides* | $F(2,4)=0.03$ $p\|p_{perm}=.96$ | $F(2,4)=1.59$ $p\|p_{perm}=.21$ | $F(2,4)=0.8$ $p\|p_{perm}=.45$ | $F(2,4)=0.13$ $p\|p_{perm}=.87$ | $F(2,4)=1.49$ $p\|p_{perm}=.25$ | $F(2,4)=0.71$ $p\|p_{perm}=.51$ |
| ***Interaction*** | | | | | | |
| *Stimuli * Species* | $F(1,2)=1.15$ $p\|p_{perm}=.74$ | $F(1,2)=2.23$ $p\|p_{perm}=.15$ | $F(1,2)=1.25$ $p\|p_{perm}=.27$ | $F(1,2)=0.05$ $p\|p_{perm}=.82$ | $F(1,2)=0.01$ $p\|p_{perm}=1$ | $F(1,2)=4.13$ $p\|p_{perm}=.05$ |
| *Stimuli * Channels* | $F(2,3)=1.3$ $p\|p_{perm}=.29$ | $F(2,3)=0.17$ $p\|p_{perm}=.85$ | $F(2,3)=0.26$ $p\|p_{perm}=.77$ | $F(2,3)=0.03$ $p\|p_{perm}=.96$ | $F(2,3)=0.09$ $p\|p_{perm}=.92$ | $F(2,3)=0.02$ $p\|p_{perm}=.98$ |
| *Species * Channels* | $F(2,3)=0.35$ $p\|p_{perm}=.72$ | $F(2,3)=0.07$ $p\|p_{perm}=.94$ | $F(2,3)=0.01$ $p\|p_{perm}=1$ | $F(2,3)=0.01$ $p\|p_{perm}=.99$ | $F(2,3)=0.06$ $p\|p_{perm}=.96$ | $F(2,3)=0.01$ $p\|p_{perm}=.99$ |
| *Stimuli * Species * Channels* | $F(2,3)=0.65$ $p\|p_{perm}=.53$ | $F(2,3)=0.18$ $p\|p_{perm}=.83$ | $F(2,3)=0.25$ $p\|p_{perm}=.78$ | $F(2,3)=0.04$ $p\|p_{perm}=.96$ | $F(2,3)=0.05$ $p\|p_{perm}=.95$ | $F(2,3)=0.05$ $p\|p_{perm}=.95$ |

**Table S2:** Results from permutation tests with 2000 permutations for each channel and subject in left hemisphere. Abbreviations: (ch1) channel 1; (ch2) channel 2; (ch3) channel 3. Analyses toward significance are highlighted in light green.

| | Talma | | | Rubis | | | Chet | | |
|---|---|---|---|---|---|---|---|---|---|
| | **ch1** | **ch2** | **ch3** | **ch1** | **ch2** | **ch3** | **ch1** | **ch2** | **ch3** |
| ***Main effect*** | | | | | | | | | |
| *Species* | $F(1,2)=4.07$ $p\|p_{perm}=.05$ | $F(1,2)=4.07$ $p=.05$ $p_{perm}=.04$ | $F(1,2)=4.07$ $p\|p_{perm}=.05$ | $F(1,2)=2.32$ $p\|p_{perm}=.13$ | $F(1,2)=2.32$ $p\|p_{perm}=.13$ | $F(1,2)=2.32$ $p\|p_{perm}=.13$ | $F(1,2)=0.22$ $p=.65$ $p_{perm}=.63$ | $F(1,2)=0.22$ $p=.65$ $p_{perm}=.64$ | $F(1,2)=0.22$ $p\|p_{perm}=.65$ |
| *Stimuli* | $F(1,2)=1.8$ $p\|p_{perm}=.19$ | $F(1,2)=1.8$ $p\|p_{perm}=.18$ | $F(1,2)=1.8$ $p\|p_{perm}=.19$ | $F(1,2)=0.22$ $p=.64$ $p_{perm}=.65$ | $F(1,2)=0.22$ $p\|p_{perm}=.64$ | $F(1,2)=0.22$ $p\|p_{perm}=.64$ | $F(1,2)=0.69$ $p\|p_{perm}=.41$ | $F(1,2)=0.69$ $p=.41$ $p_{perm}=.4$ | $F(1,2)=0.69$ $p=.41$ $p_{perm}=.42$ |
| *Sides* | $F(2,4)=1.4$ $p\|p_{perm}=.26$ | $F(2,4)=1.4$ $p\|p_{perm}=.26$ | $F(2,4)=1.4$ $p\|p_{perm}=.26$ | $F(2,4)=0.12$ $p=.89$ $p_{perm}=.88$ | $F(2,4)=0.12$ $p=.89$ $p_{perm}=.88$ | $F(2,4)=0.12$ $p\|p_{perm}=.89$ | $F(2,4)=0.42$ $p\|p_{perm}=.66$ | $F(2,4)=0.42$ $p=.66$ $p_{perm}=.65$ | $F(2,4)=0.42$ $p=.66$ $p_{perm}=.65$ |
| ***Interaction*** | | | | | | | | | |
| *Species* Stimuli* | $F(1,2)=2.15$ $p=.15$ $p_{perm}=.16$ | $F(1,2)=2.15$ $p\|p_{perm}=.15$ | $F(1,2)=2.15$ $p=.15$ $p_{perm}=.16$ | $F(1,2)=0.05$ $p\|p_{perm}=.82$ | $F(1,2)=0.05$ $p\|p_{perm}=.82$ | $F(1,2)=0.05$ $p\|p_{perm}=.82$ | $F(1,2)=3.75$ $p=.07$ $p_{perm}=.06$ | $F(1,2)=3.75$ $p\|p_{perm}=.07$ | $F(1,2)=3.75$ $p=.07$ $p_{perm}=.06$ |

**Table S3:** Results from permutation tests with 2000 permutations for each channel and subject in right hemisphere. Abbreviations: (ch1) channel 1; (ch2) channel 2; (ch3) channel 3. Analyses toward significance are highlighted in light green.

| | **Talma** | | | **Rubis** | | | **Chet** | | |
|---|---|---|---|---|---|---|---|---|---|
| | **ch1** | **ch2** | **ch3** | **ch1** | **ch2** | **ch3** | **ch1** | **ch2** | **ch3** |
| ***Main effect*** | | | | | | | | | |
| *Species* | $F(1,2)=0.15$ $p\|p_{perm}=.70$ | $F(1,2)=0.15$ $p\|p_{perm}=.69$ | $F(1,2)=0.15$ $p\|p_{perm}=.69$ | $F(1,2)=0.12$ $p\|p_{perm}=.72$ | $F(1,2)=0.12$ $p=.72$ $p_{perm}=.73$ | $F(1,2)=0.12$ $p\|p_{perm}=.72$ | $F(1,2)=3.74$ $p\|p_{perm}=.07$ | $F(1,2)=3.74$ $p=.07$ $p_{perm}=.06$ | $F(1,2)=3.74$ $p=.07$ $p_{perm}=.06$ |
| *Stimuli* | $F(1,2)=0.03$ $p=.98$ $p_{perm}=.99$ | $F(1,2)=0.03$ $p=.98$ $p_{perm}=.99$ | $F(1,2)=0.03$ $p=.98$ $p_{perm}=.99$ | $F(1,2)=1.12$ $p=.29$ $p_{perm}=.3$ | $F(1,2)=1.12$ $p\|p_{perm}=.29$ | $F(1,2)=1.12$ $p=.29$ $p_{perm}=.28$ | $F(1,2)=0.09$ $p=.76$ $p_{perm}=.75$ | $F(1,2)=0.09$ $p\|p_{perm}=.76$ | $F(1,2)=0.09$ $p\|p_{perm}=.76$ |
| *Sides* | $F(2,4)=0.04$ $p\|p_{perm}=.96$ | $F(2,4)=0.04$ $p\|p_{perm}=.96$ | $F(2,4)=0.04$ $p\|p_{perm}=.96$ | $F(2,4)=0.75$ $p=.48$ $p_{perm}=.47$ | $F(2,4)=0.75$ $p=.48$ $p_{perm}=.49$ | $F(2,4)=0.75$ $p=.48$ $p_{perm}=.47$ | $F(2,4)=0.91$ $p=.42$ $p_{perm}=.4$ | $F(2,4)=0.91$ $p=.42$ $p_{perm}=.43$ | $F(2,4)=0.91$ $p\|p_{perm}=.42$ |
| ***Interaction*** | | | | | | | | | |
| *Species* Stimuli* | $F(1,2)=0.11$ $p\|p_{perm}=.74$ | $F(1,2)=0.11$ $p=.74$ $p_{perm}=.73$ | $F(1,2)=0.11$ $p=.74$ $p_{perm}=.75$ | $F(1,2)=1.22$ $p=.27$ $p_{perm}=.28$ | $F(1,2)=1.22$ $p\|p_{perm}=.27$ | $F(1,2)=1.22$ $p\|p_{perm}=.27$ | $F(1,2)=0.01$ $p\|p_{perm}=1$ | $F(1,2)=0.01$ $p\|p_{perm}=1$ | $F(1,2)=0.01$ $p\|p_{perm}=1$ |
